# Tensor Decomposition-based Unsupervised Feature Extraction Succeeded in Identification of Differentially Expressed Transcripts from Redundant *de novo* Transcriptome of *Planarian*

**DOI:** 10.1101/2021.06.15.448531

**Authors:** Makoto Kashima, Nobuyoshi Kumagai, Hiromi Hirata, Y-h. Taguchi

**Author notes:** Department of Biomolecular Science, Faculty of Science, Toho University, Miyama 2-2-1, Funabashi, 274-8510, Chiba, Japan (Makoto Kashima).

## Abstract

RNA-Seq data analysis of non-model organisms is often difficult because of the lack of a well-annotated genome. However, in non-model organisms, contigs can be generated by *de novo* assembling. This can result in a large number of transcripts, making it difficult to easily remove redundancy. A large number of transcripts can also lead to difficulty in the recognition of differentially expressed transcripts (DETs) between more than two experimental conditions, because *P*-values must be corrected by considering multiple comparison corrections whose effect is enhanced as the number of transcripts increases. Heavily corrected *P*-values often fail to take sufficiently small *P*-values as significant. In this study, we applied a recently proposed tensor decomposition (TD)-based unsupervised feature extraction (FE) to the RNA-seq data obtained for a non-model organism, planarian *Dugesia japonica*; Although we used de novo assembled transcriptome reference with high redundancy, we successfully obtained a larger number of transcripts whose expression was altered between normal and defective samples as well as during time development than those identified by a conventional method. TD-based unsupervised FE is expected to be an effective tool that can identify a substantial number of DETs, even when a poorly annotated genome is available.

## 1. Introduction

Identification of differentially expressed transcripts (DETs) [1,2] between more than two distinct experimental conditions is the starting point of RNA-seq data analysis, as DETs are expected to be related to the experimental conditions considered, for example, diseases. Once DETs are successfully identified, it is possible to study the biological properties that are enriched in a set through a statistical test such as gene ontology enrichment analysis [3]. Thus, unless DETs can be successfully identified, it is very difficult to make use of RNA-seq data to identify the biological processes that take place during experiments using RNA-seq datasets.

DET identification in model organisms has been well-established. Once short reads are successfully mapped to the genome, we can concentrate those mapped to well-annotated regions; for example, protein-coding genes. Then, the number of reads can be summed up within the considered regions, for example, the gene body, then it is possible to estimate the amount of expression of individual genes [4]. Another way is that reference transcriptome sequences are generated based on gene models [2,5], following by mapping short reads onto it. Then, quantification is performed considering information of isoform and reads mapped onto several reference sequences [2,6–8]. The latter approaches tend to be superior in quantification accuracy than the former [9]. In contrast, DET identification of non-model organisms is not straightforward. Because of the lack of a well-annotated genome, no short reads can be mapped to it. Instead of the genome, it is possible to assemble the contigs by assembling *de novo*, and short reads can then be mapped toward these contigs. However, there is one problem with this strategy; it is never possible to be confident that the parts of the assembled contigs are unique. Some contigs may overlap with some parts of another contig [10,11]. This results in so-called multiple mapping, which drastically reduces the accuracy of the estimated amount of transcript [12]. In addition, the number of contigs can often be far larger than the true number of transcripts because of the above-mentioned redundancy of contigs. e.g. the number of contigs can often be in millions. This causes serious problems of drastic reduction of statistical power in DET identification. In popular DET identification methods such as DESeq2 [13], the statistical tests check how significant the observed distinct expression of a transcript is under the null hypothesis and calculate *P*-values, which must be corrected by considering multiple comparison corrections. As the number of multiple comparisons is equivalent to the number of contigs, too many contigs (e.g. millions) results in false negative identification of DETs.

To address this problem, we applied tensor decomposition (TD)-based unsupervised feature extraction (FE) [14] to the RNA-seq data. As proof of concept, here, we analyzed RNA-Seq data of a planarian *Dugesia japonica*, one of whose *de novo* trascriptome assembly contains as many as 2.8 *×* 105 [15], which is much larger than the expected number of true genes of the planarian [16]. TD based unsupervised FE could successfully select a larger number of contigs whose expression was altered during time development as well as was distinct between normal and defective samples than a conventional method DESeq2. This suggests the usefulness of TD-based unsupervised FE when it is applied to RNA-seq data analysis of non-model organisms, from which too many redundant contigs are often obtained.

## 2. Materials and Methods

### 2.1 Maintenance of planarian

A clonal strain of planarian *D. japonica*, a sexualizing super planarian (2n = 16), was used in this study [17]. The planarians were fed edible chicken liver once or twice a week. The planarians that were approximately 5 mm in length were starved for at least one week prior to the following experiments.

### 2.2 Double-stranded RNA (dsRNA) synthesis

dsRNA was synthesised as described by [18]. We prepared templates flanked by the T7 promoter for dsRNA synthesis by using polymerase chain reaction (PCR) for each EST (*Djpi-wiA* [*Dj_aH_000_03609HH*], *DjpiwiB* [*Dj_aH_221_M14*], *DjpiwiC* [*Dj_aH_000_05977HH*], *Djhp1-1* [*Dj_aH_000_01636HH*], *Djima1-1* [*Dj_aH_313_F03*]). The primers for the PCR reaction were as follows (5’ to 3’): Zap Linker + T7: GATCACTAATACGACTCAC-TATAGGGGAATTCGGCACGAGG M13 Rev: GTTTTCCCAGTCACGACGTTGTAA.

### 2.3 Feeding RNA interference

RNA interference was conducted as described previously [18]. dsRNA-containing food consisting of 25 *µ*L of chicken liver solution, 6 *µ*L of 2% agarose, and 6.5 *µ*L of 2.0 *µ*g/*µ*L dsRNA was synthesised in vitro. We fed the planarians twice at intervals of two days. Control animals were fed food containing eGFP dsRNA. Six individual planarians were sacrificed every day from day 1 to 16 after the second feeding.

### 2.4 Total RNA purification

Total RNA was extracted from each individual planarian using the “Direct-TRI” method [19]. Briefly, a planarian was lysed using TRI Reagent-LS (Molecular Research Center, Cincinnati, OH, USA). Then, the lysate was purified using an AcroPrep Advance 96-well long tip filter plate for nucleic acid binding (Pall, Port Washington, NY, USA). RNA was eluted with 30 µL nuclease-free water.

### 2.5 RNA-Seq library preparation and sequencing

3’ mRNA-Seq was conducted using the Lasy-Seq ver. 1.1 Protocol (https://sites.google.com/view/lasy-seq/) [20,21]. Briefly, 100 ng of total RNA was reverse transcribed using an RT primer with an index and SuperScript IV reverse transcriptase (Thermo Fisher Scientific, Waltham, MA, USA). All RT mixtures of the samples were pooled and purified using an equal volume of AMpure XP beads (Beckman Coulter, Brea, CA, USA) according to the manufacturer’s instructions. Second-strand synthesis was conducted on the pooled samples using RNaseH (5 U/*µ*L, Enzymatics, Beverly, MA, USA), and DNA polymerase I (10 U/*µ*L, Enzymatics, Beverly, MA, USA). To avoid carryover of large amounts of rRNAs, the mixture was subjected to RNase treatment using RNase T1 (Thermo Fisher Scientific, Waltham, MA, USA). Then, purification was conducted with a 0.8× volume of AMpure XP beads. Fragmentation, end-repair, and A-tailing were conducted using a 5× WGS fragmentation mix (Enzymatics, Beverly, MA, USA). The adapter for Lasy-Seq was ligated using 5× Ligation Mix (Enzymatics, Beverly, MA, USA), and the adapter-ligated DNA was purified twice with a 0.8× volume of AMpure XP beads. After the optimisation of PCR cycles for library amplification by qPCR using Evagreen, 20× in water (Biotium, Fremont, CA, USA) and the QuantStudio5 Real-Time PCR System (Applied Biosystems, Waltham, MA, USA), the library was amplified using KAPA HiFi HotStart ReadyMix (KAPA BIOSYSTEMS, Wilmington, MA, USA) on the ProFlex PCR System (Applied Biosystems, Waltham, MA, USA). The amplified library was purified using an equal volume of AMpure XP beads. One microliter of the library was then used for electrophoresis using a Bioanalyzer 2100 with the Agilent High Sensitivity DNA kit (Agilent Technologies, Santa Clara, CA, USA) to check for quality. Sequencing of 150 bp paired-end reads was performed using HiSeq X Ten (Illumina, San Diego, CA, USA).

### 2.6 Mapping and gene quantification

Read 1 reads were processed with fastp (version 0.21.0) [22] using the following parameters: trim_poly_x −w 20 –adapter_sequence=AGATCGGAAGAGCACACGTCTGAACTCCAGTCA –adapter_sequence_r2= AGATCGGAAGAGCGTCGTGTAGGGAAAGAGTGT-l 31. The trimmed reads were then mapped to a *D. japonica* reference sequence of GJEZ00000000 in the TSA repository, which is a de novo transcriptome assembly generated in a previous study [15], using BWA mem (version 0.7.17-r1188) [23] with the default parameters. The read count for each gene was calculated with salmon using −l IU, which specifies the library type (version v0.12.0) [6]. All quantification results and sample information of the RNA-seq analysis were deposited as GSE174855 in the GEO repository.

### 2.7 Mapping to oxford genome

To see if the analysis of contig without genome is reasonable, we mapped the obtained short reads to genome, too. The reference transcriptome sequences were generated based on a draft genome [24] and gene models [25] by using gffread (v0.12.1) [5]. Then, mapping and quantification were conducted as described above.

### 2.8 Comparison of the de novo transcriptome assembly and oxford references

BLASTN anaysis (version 2.9.0+) [26] was performed using the oxford references as query and the de novo transcriptome assembly as reference. Best hit of each oxford reference sequence was refer to the corresponding contig in the de novo transcriptome assembly.

### 2.9 TD-based unsupervised FE

The number of reads mapped to individual contigs was formatted as a tensor, *x*_*ijtk*_ *∈ ℝ*^*N×*6*×T×*6^, which represents the expression of the *i*th contig (*N* = 278, 167) or gene (*N* = 24, 090) of the *k*th biological replicates at the *t*th day after the *j*th RNAi was performed. The 14th dataset for contig was excluded because it included serious outliers. Thus, *t ≤* 13 corresponds to *t* days after the treatment, whereas *t ≥* 15 corresponds to *t* + 1 days after the treatment. Then *T* = 15 for contig. Since it was not a case for genes, *T* = 16.

*x*_*ijtk*_ is normalised such that ∑_*i*_ *x*_*ijtk*_ = 0 and 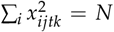. Then, we applied higher order singular value decomposition (HOSVD) [14] and got

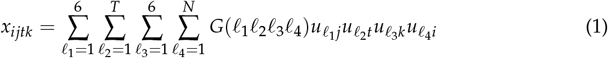

where *G ∈* ℝ^6*×T×*6*×N*^ is the core tensor that represents the weight of the product 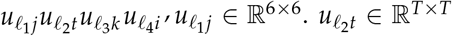 and 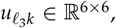 and 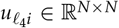 are singular value matrices that are orthogonal matrices.

First, we needed to identify 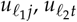, and 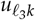 that satisfy

- 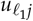 that exhibits distinction between normal and defective samples.
- 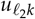 with time (days) dependence.
- 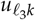 independent of individual biological replicates, *k*, i.e. with constant values.

After identifying *ℓ*_1_, *ℓ*_2_, and *ℓ*_3_ that satisfy the above requirements, we sought a set of *ℓ*_4_s, 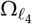, associated with the larger |*G*(*ℓ*_1_*ℓ*_2_*ℓ*_3_*ℓ*_4_)|s given *ℓ*_1_, *ℓ*_2_, and *ℓ*_3_, because 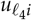 s are expected to have larger absolute values for the genes whose expression is associated with the above requirement. Then, we attributed *P*-values to the *i*th contig by assuming that 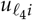 obeys a multiple Gaussian distribution (null hypothesis)

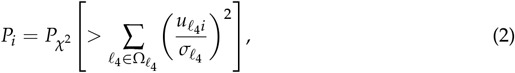

where 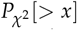 is the cumulative probability distribution of the *χ*^2^ distribution when the argument takes larger values than *x*. The obtained *P*_*i*_s are corrected by the BH criterion [14], and contigs associated with an adjusted *P*_*i*_ less than 0.01 were selected.

### 2.10 DESeq2

DESeq2 [13] (version 1.30.0) was applied to either comparison between two groups of RNAi treatments, that is, *gfp, DjpiwiA* and others (*Djhp1-1, Djima1-1, DjpiwiB, DjpiwiC*) or that between two time periods, that is, *t* = 1, 2, *• • •*, 8, and others (*t* = 9, 10, 11, 12, 13, 15, 16). We selected genes associated with adjusted *P*-values less than 0.01, which is the same as that used for TD-based unsupervised FE. All other parameters were taken as the default values.

### 2.11 GO term enrichment analysis

BLASTX (version 2.9.0+) [26] was used for annotation of planarian contigs/genes. All mouse protein data (GRCm3.8) [27] was used as BLAST subject sequences. Best hit of each transcript was used for the annotation of planarian transcripts. Then, GO term annotation to planarian transcripts was conducted by using biomaRt (version 2.46.0) [28],GO.db (version 3.12.1) [29], and org.Mm.eg.db (version 3.11.4) [30]. GO term enrichment analysis was conducted with a function in R: fisher.test. Then, the adjustment for multiple comparisons against *P*-values was performed using the Benjamini–Hochberg (BH) method [31] using the p.adjust function in R. Finally, we listed only GO terms of the most offspring with the adjusted P-value *<* 0.05.

## 3. Results

The RNA-Seq data were derived from RNAi experiments designed to identify transcripts related to regenerative defects caused by knockdown of *DjpiwiB* and *DjpiwiC* in a planarian *Dugesia japonica*. In addition to a control and three planarian *piwi* genes’, we conducted RNAi experiments against two planarian genes, *Djhp1-1* and *Djima1-1*, because they are homologous genes in fruit fly are known to be partners of PIWI proteins [32,33]. The each targeted gene was significantly knocked-down under each RNAi treatment (Fig. 1).

**Figure 1.**
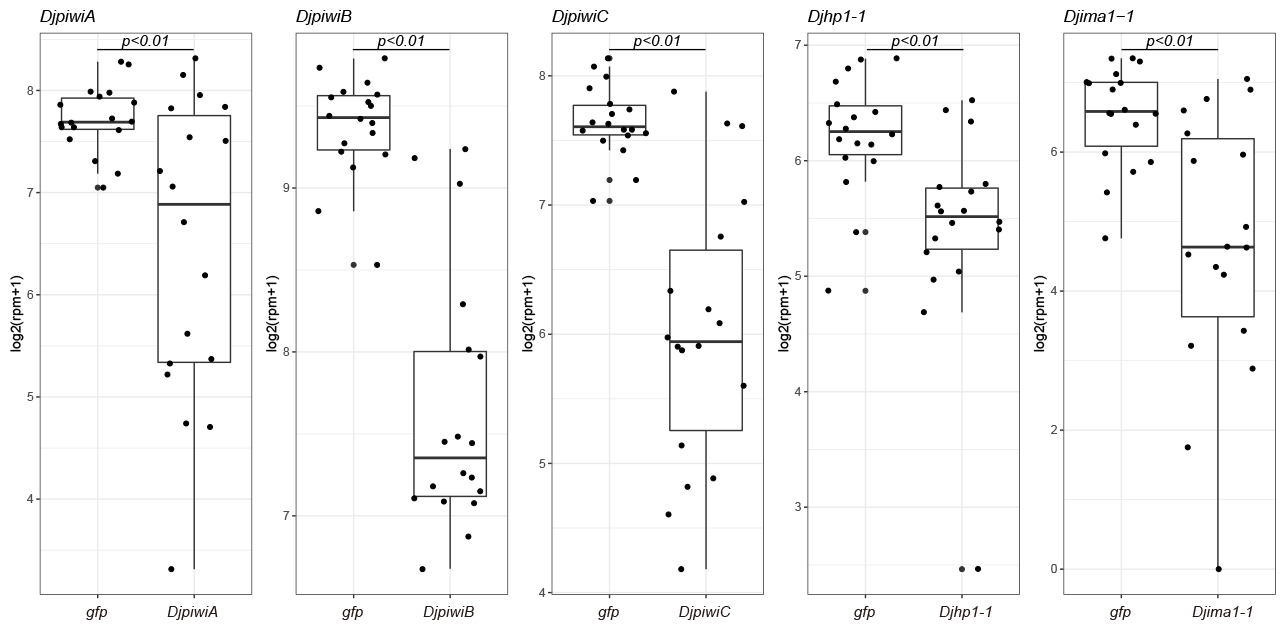
log2(rpm+1) of RNAi target genes at 3, 4, and 5 days into the RNAi treatment. P-value were calculated with t-test.

First, we attempted to identify which *u*_*ℓ*1 *j*_ is associated with the desired property. Figure 2 shows *u*_2*j*_ that represents distinction between the RNAi treatments. The phenotype of RNAi of *Djhp1-1, Djima1-1, DjpiwiB*, and *DjpiwiC* is distinct from that of *gfp* and *DjpiwiA*. While *gfp* is the control treatment and KD of *piwiA* is known not to affect (inhibit) the regeneration of planarians, RNAi of *Djhp1-1, Djima1-1, DjpiwiB*, and *DjpiwiC* causes regenerative defects [15,34,35]. Thus, *u*_2*j*_ coincides with the distinction between normal and defective samples.

**Figure 2.**
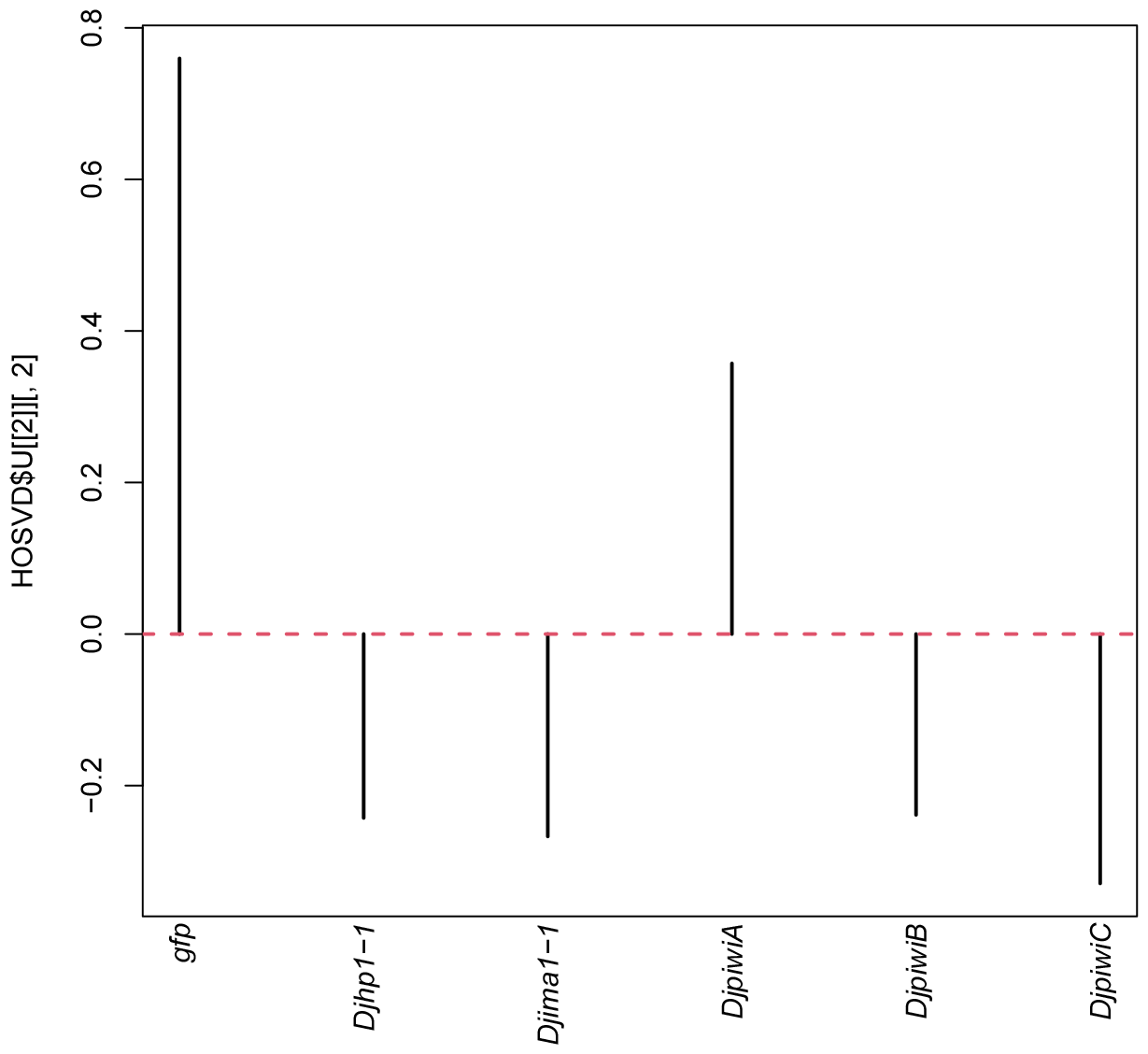
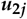 that represent 2nd singular value vectors attributed to RNAi experiments that coincides with the distinction between normal and defective samples. That is, *gfp* and KD of *piwiA* is known not to affect (inhibit) the regeneration of planarians and RNAi of *Djhp1-1, Djima1-1, DjpiwiB*, and *DjpiwiC* causes regenerative defects [15,34,35].

Next, we attempted to identify which 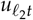 coincides with time development. Figure 3 shows *u*_2*t*_ that exhibits the desired time development. Several days after RNAi treatment, RNAi appeared to have an effect, which then gradually decreased. The previous study showed *DjpiwiB*-knocked-down planarians lose their regenerative ability at seven days after the RNAi treatment [15]. This time point corresponds to the first date when *u*_2*t*_ takes positive values (the sign of *u*_2*t*_ does not have any meaning because only the product of 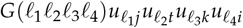 matters, not individual terms in the product (see eq.(1)). Thus, this time development seemed to coincide with the experimental procedures. The reason why *u*_2*t*_ takes almost zero might be because it requires several days until RNAi starts to affect the transcriptome.

**Figure 3.**
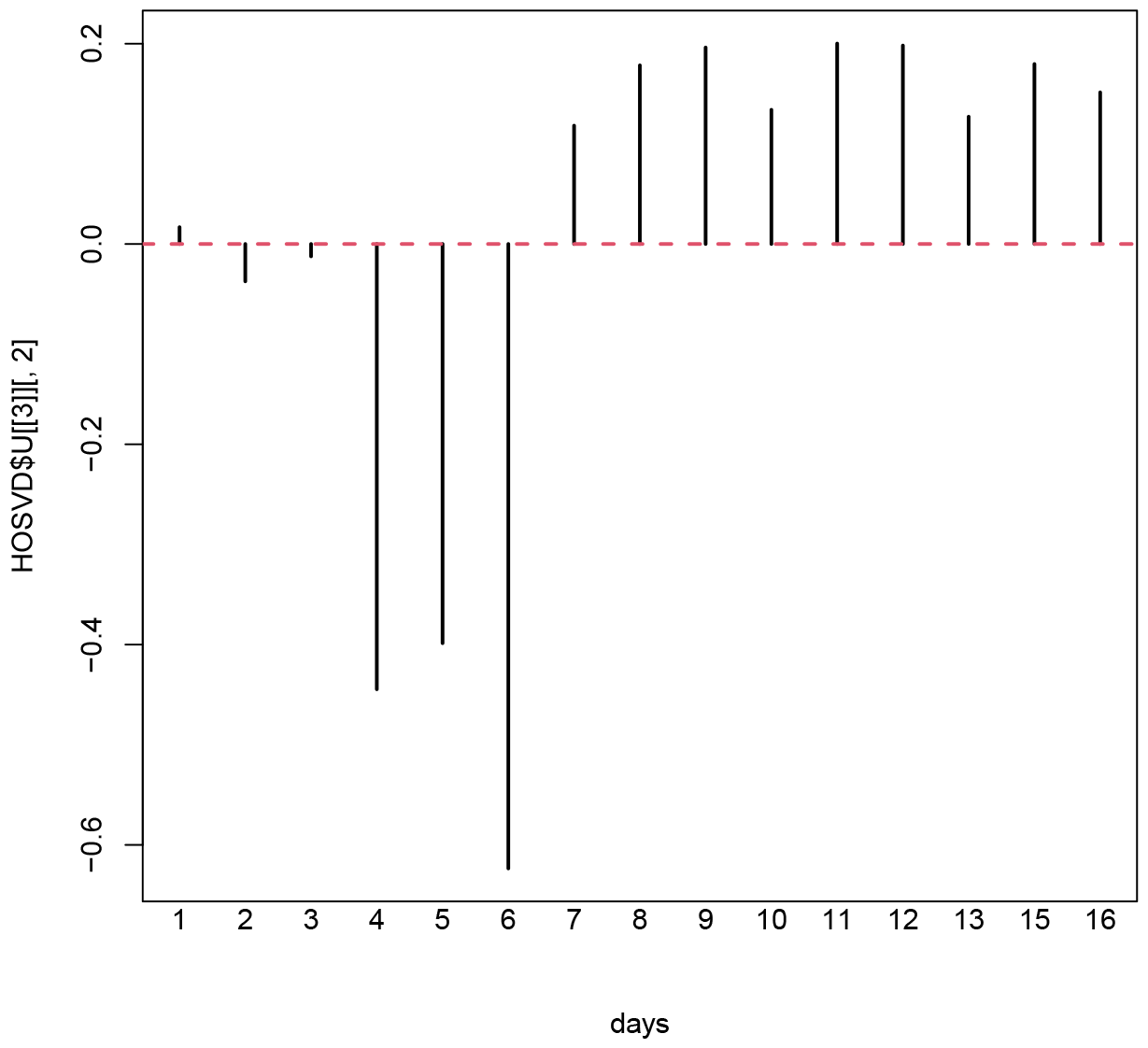
***u***_**2*t***_ that represent 2nd singular value vectors attributed to days after the treatments. The time point where sign of *u*_2*t*_ changes coincides with time point when regenerative defects of *DjpiwiB* RNAi appears.

We also noticed that *u*_1*k*_ has almost constant values independent of the biological replicate, *k* (Fig. 4); this suggests that the expression of genes associated with *u*_1*k*_ is likely to be common among the six biological replicates.

**Figure 4.**
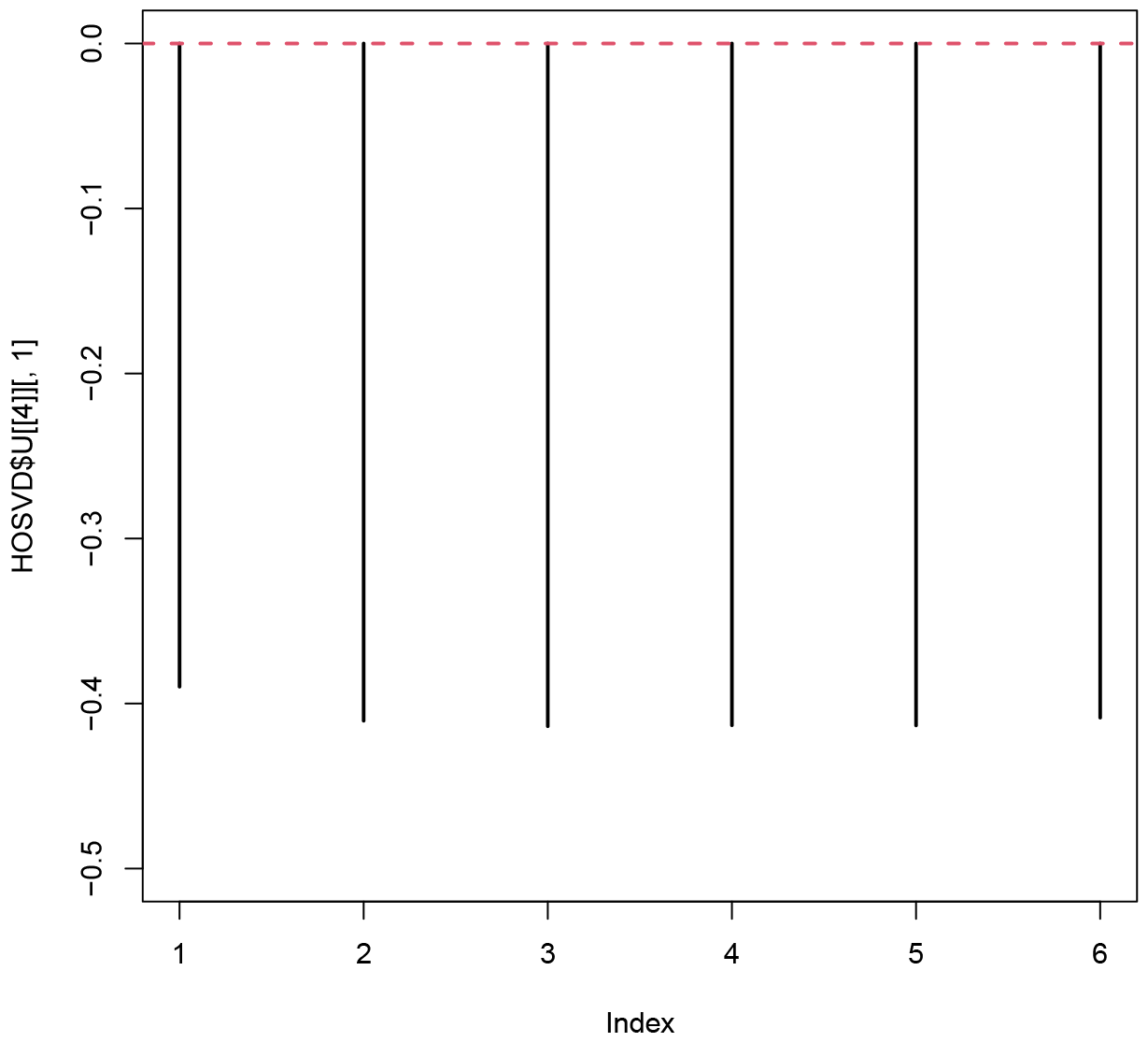
***u***_**1*k***_ that represent 1st singular value vectors attributed to biological replicates. The values indicate there is no bias between the biological replicates.

We then determine which *ℓ*_4_ had the largest |*G*(2, 2, 1, *ℓ*_4_)|. Table 1 shows *G*(2, 2, 1, *ℓ*_4_). *ℓ*_4_ = 1, 3, 4 had a larger |*G*| value. It is not possible to employ all three *ℓ*_4_s to compute *P*_*i*_ because |*G*(1, 1, 1, 1)| is much larger than *G*(2, 2, 1, 1) (not shown here). This means that *u*_1*i*_ is more coincident with *ℓ*_1_ = *ℓ*_2_ = *ℓ*_3_ = 1, which we are not interested in. Thus, we decided to employ *ℓ*_4_ = 3, 4 to attribute *P*-values to the *i*th contig (that is, 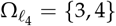 in Eq.(2)). *P*_*i*_s are corrected as described in the Methods section, and contigs with adjusted *P*_*i*_ less than 0.01 were selected.

**Table 1.** *G*(2, 2, 1, *ℓ*_4_)

| $\ell_1$ | $G(2, 2, 1, \ell_4)$ |
| --- | --- |
| 1 | 227.87614 |
| 2 | -8.36258 |
| 3 | -113.10063 |
| 4 | 250.20079 |
| 5 | -43.37827 |
| 6 | -64.02292 |
| 7 | 37.49047 |
| 8 | 12.84754 |
| 9 | 31.20391 |
| 10 | -21.70284 |

After applying TD-based unsupervised FE to RNA-seq data obtained from Planarian as described in the Methods section, we obtained 155 contigs as those whose expression was altered during time development as well as distinct between normal and defective samples. To confirm whether we could successfully identify genes with altered expression, we applied statistical tests that validated significant dependence upon time as well as significant distinction between normal and defective samples. First, we applied the *t* test to determine whether gene expression was distinct between *t ≤* 8 and *t ≥* 9 (Table 2). There were at least non-zero genes expressed differently between *t ≤* 8 and *t ≥* 9.

**Table 2.** The number of contigs identified by the *t* test to be distinct between *t ≤* 8 and *t ≥* 9.

| RNAi | netative | positive |
| --- | --- | --- |
| *gfp* | 89 | 66 |
| *Djhp1-1* | 96 | 59 |
| *Djima1-1* | 139 | 16 |
| *DjpiwiA* | 89 | 66 |
| *DjpiwiB* | 102 | 53 |
| *DjpiwiC* | 151 | 4 |

We also checked whether genes were expressed differently on individual days between normal and defective samples (Table 3). In this case, more than half of the genes were differentially expressed between normal and defective samples.

**Table 3.** The number of contigs identified by the *t* test to be distinct between normal and defective samples.

| days | negative | positive |
| --- | --- | --- |
| 1 | 76 | 79 |
| 2 | 118 | 37 |
| 3 | 41 | 114 |
| 4 | 69 | 86 |
| 5 | 78 | 77 |
| 6 | 90 | 65 |
| 7 | 48 | 107 |
| 8 | 73 | 82 |
| 9 | 125 | 30 |
| 10 | 93 | 62 |
| 11 | 65 | 90 |
| 12 | 46 | 109 |
| 13 | 58 | 97 |
| 15 | 95 | 60 |
| 16 | 50 | 105 |

To compare the statistical power of TD-based unsupervised FE and conventional methods, a conventional method DESeq2 [13] was applied to either comparison between two groups of RNAi treatments, that is, *gfp, DjpiwiA* and others (*Djhp1-1, Djima1-1, DjpiwiB, DjpiwiC*) or that between two time periods, that is, *t* = 1, 2, *…*, 8, and others (*t* = 9, 10, 11, 12, 13, 15, 16). Although we selected genes associated with adjusted *P*-values less than 0.01, which is the same as that used for TD-based unsupervised FE, DESeq2 identified as few as 10 and 5 contigs for those expressed differently between normal and defective samples, respectively.

Furthermore, in order to evaluate 155 contigs from the viewpoints of biological functions, we performed a Basic Local Alignment Search Tool (BLAST) match between 155 contigs and a well-annotated mouse protein database [27], followed by GO terms enrichment analysis. As a result, 14 GO terms of “Biological process”, 10 GO terms of “Molecular function”, and 4 GO terms of “Cellular component” were significantly enriched (FDR=0.05) (Fig. 5). Consistent with the enrichment of GO terms “translation” and “extracellular spac”, translation and extracellular matrix are thought to be important in planarian regeneration [36,37]. Consist with the enrichment of GO term “serine-type endopeptidase inhibitor activity”, it was reported that transcripts coding for serine-type endopeptidase inhibitors are differentially expressed during planarian regeneration [38]. Thus, we concluded that our analysis successfully selected contigs related to regenerative ability in the planarian.

**Figure 5.**
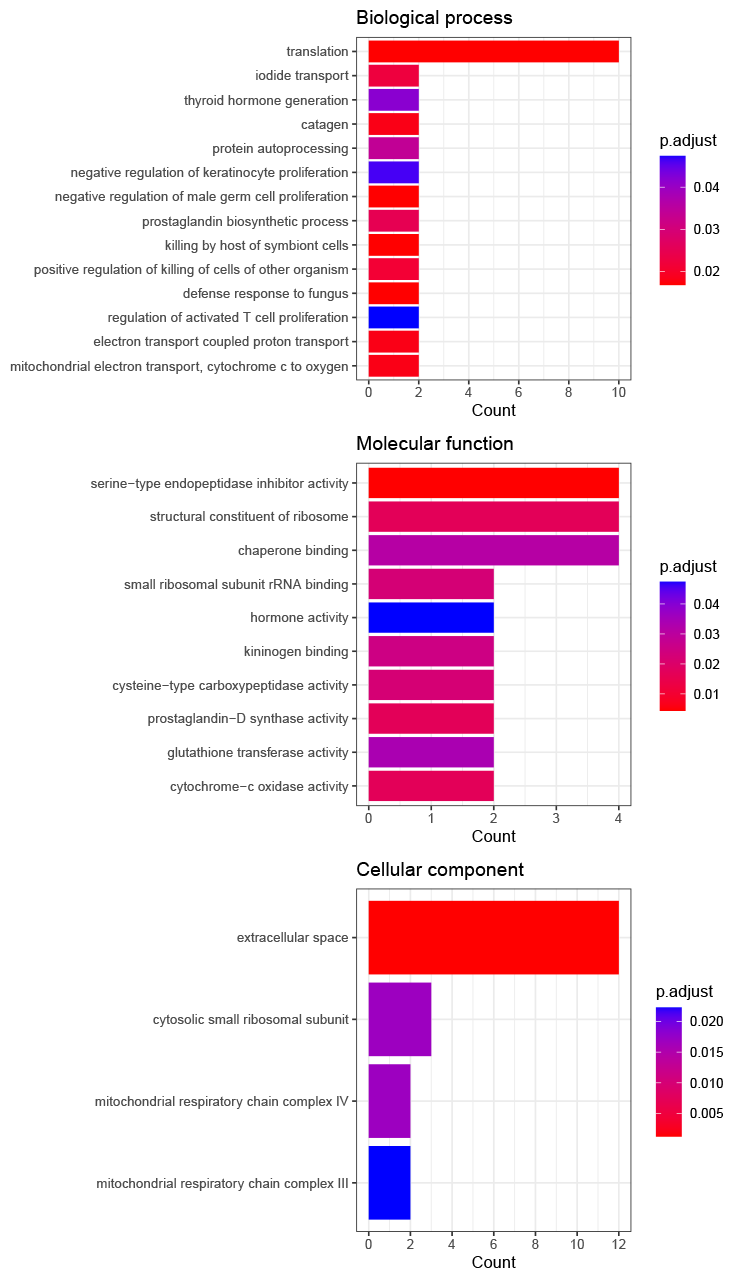
The results of GO term enrichment analysis of planarian transcripts selected by our method based on blast results against well annotated mouse proteins

## 4. Discussion

We employed unpopular TD-based unsupervised FE to identify DET despite the existence of many other conventional methods. This was to show that the problem is too difficult to resolve using other conventional methods. Therefore, we applied DESeq2 to the present dataset. As a result, we identified as few as 10 and 5 contigs for those expressed differently between normal and defective samples, respectively, and those expressed dependent upon time, and those expressed dependent upon time; these numbers are very low when compared with the number of contigs identified by TD-based unsupervised FE, 155. DESeq2 failed because there were as many as 278167 contigs, whereas the number of samples was as low as 6 *×* 15 *×* 6 = 540; its ratio was as large as 5 *×* 10^2^. It is a difficult problem to tackle using standard conventional methods designed for a much lower ratio of the number of features to the number of samples.

In order to further validate identified genes biologically, we also performed a BLAST search against nt database of 155 contigs (see Additional file 1). We found that almost half of the 155 contigs had a common match with two known long planarian transcripts; AK388828.1 and AK389113.1, and that these are likely to be alternatively spliced transcripts of the transcript. TRINITY_DN1947 (Fig. 6). Although we are unsure why Trinity [39] failed to merge these redundant contigs into one, this suggests that TD-based unsupervised FE can detect transcripts that share the same expression patterns. If individual contigs are alternative spliced short transcripts of a long transcript, which is likely to be TRINITY_DN1947, there are similar expression patterns.

**Figure 6.**
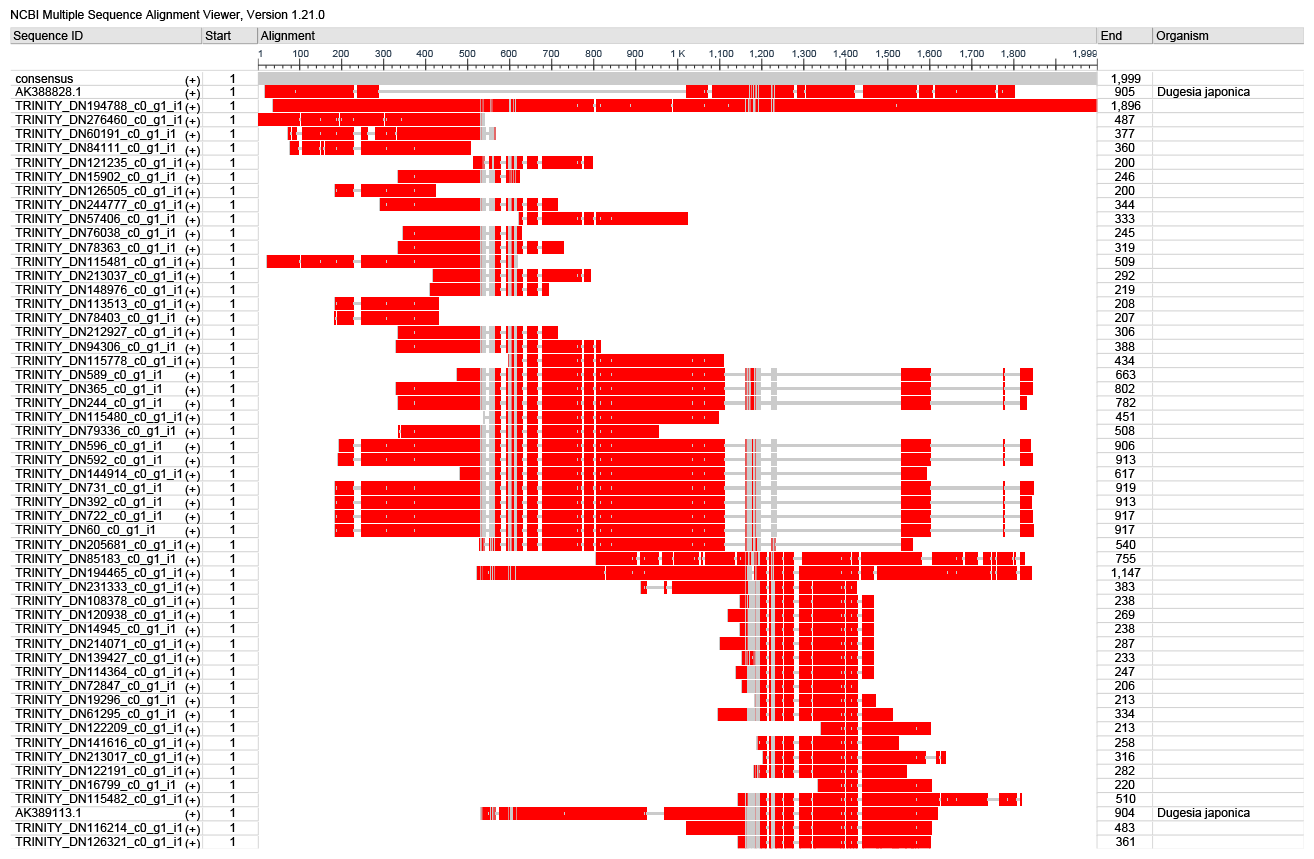
Multiple alignment of contigs. identified by using BLAST search to be aligned to known planarian transcripts, AK388828.1 or AK389113.1.

One possible objrction to the present study is that the coincidence of analyses between with and without genome and gene models. If the analyses based upon contig differ from those with genome, the present study is useless. To examine this possibility, we mapped short read to genome [24,25] and repeated the above analyses. At first, we evaluated *u*_*ℓt*_ and 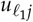 as well (Figs. 7 and 8) and we found that *u*_2*t*_ and 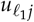, 2 *≤ ℓ*_1_ *≤* 4) coincide with the desired property; in contrast to Fig. 2 where three genes are distinct from others in *u*_2*j*_, three genes are distinct from others in 2 *≤ ℓ*_1_ *≤* 4 one by one and similar to Fig. 3, *u*_2*t*_ is dependent on *t*. Furthermore, *u*_1*k*_ is independent of *k*, replicates, as in Fig. 4 (not show here), we decided to investigate |*G*(*ℓ*_1_, 2, 1, *ℓ*_4_)|, 2 *≤ ℓ*_1_ *≤* 4 to see which *ℓ*_4_ is associated with desired properties (Fig. 9) and employed *ℓ*_4_ = 3. *P*-values are attributed to *i*s in *x*_*ijtk*_ (i.e., genes) with eq. (2) and are corrected by BH method. We identified 89 genes associated with the adjusted *P*-values less than 0.01. Since the number does not differ from that of contigs, 155, so much, analyses with and without genome seem to coincide with each other.

**Figure 7.**
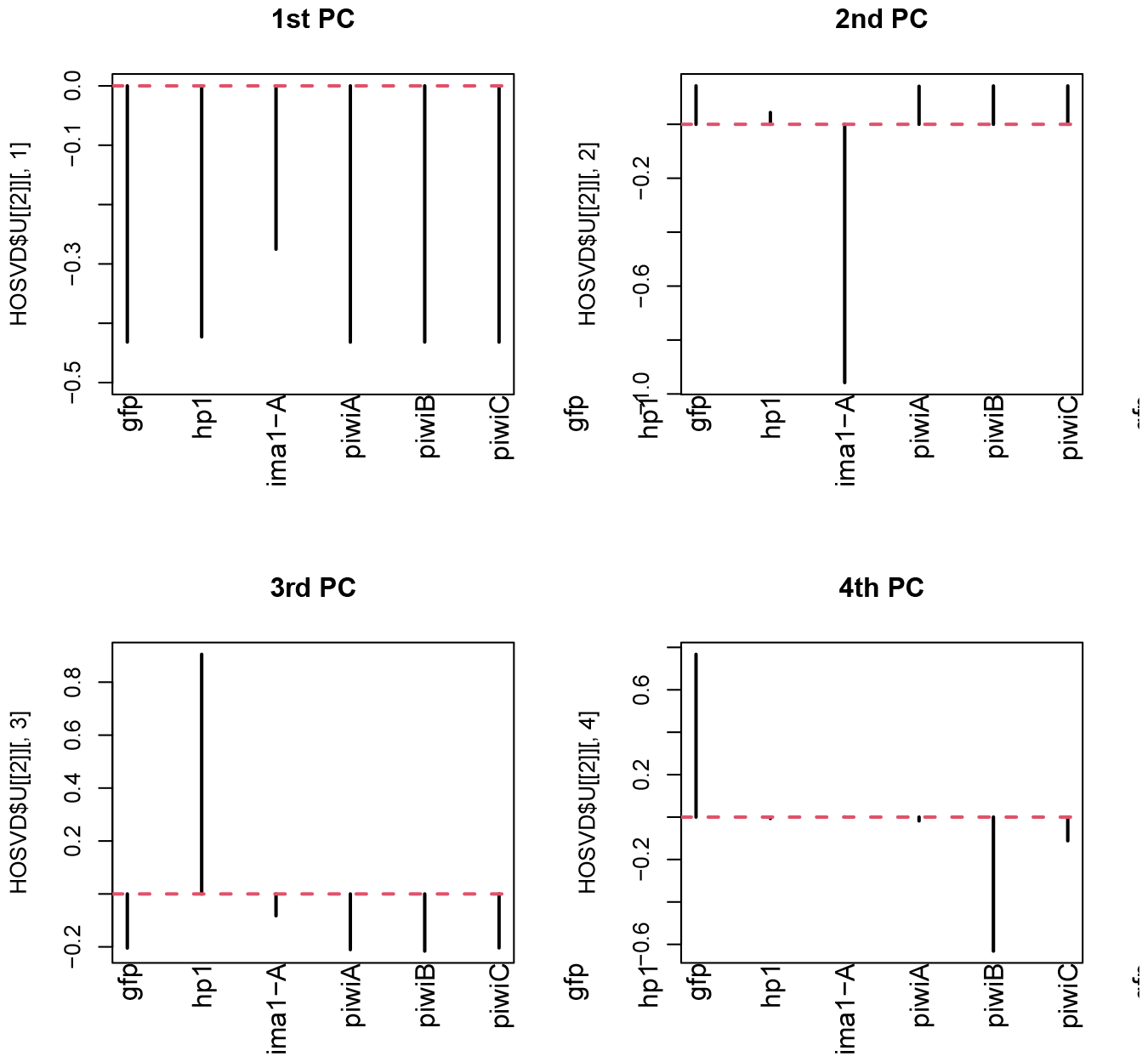
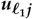 (1 *≤ ℓ*_1_ *≤* 4) that represent 1st to 4th singular value vectors attributed to RNAi experiments that coincides with the distinction between normal and defective samples.

**Figure 8.**
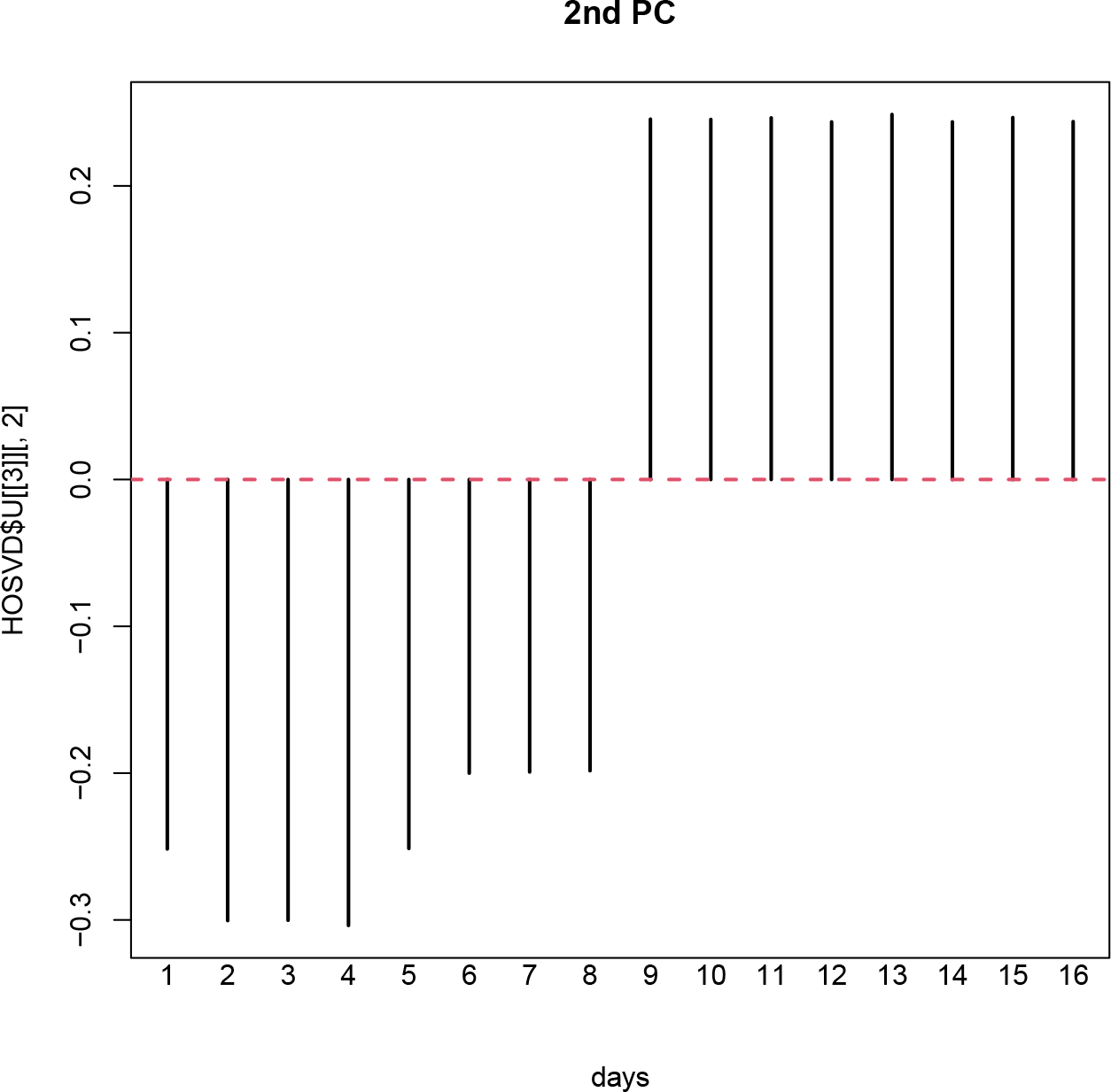
***u***_**2*t***_ that represent 2nd singular value vectors attributed to days after the treatments. The time point where sign of *u*_2***t***_ changes coincides with time point when regenerative defects of *DjpiwiB* RNAi appears.

**Figure 9.**
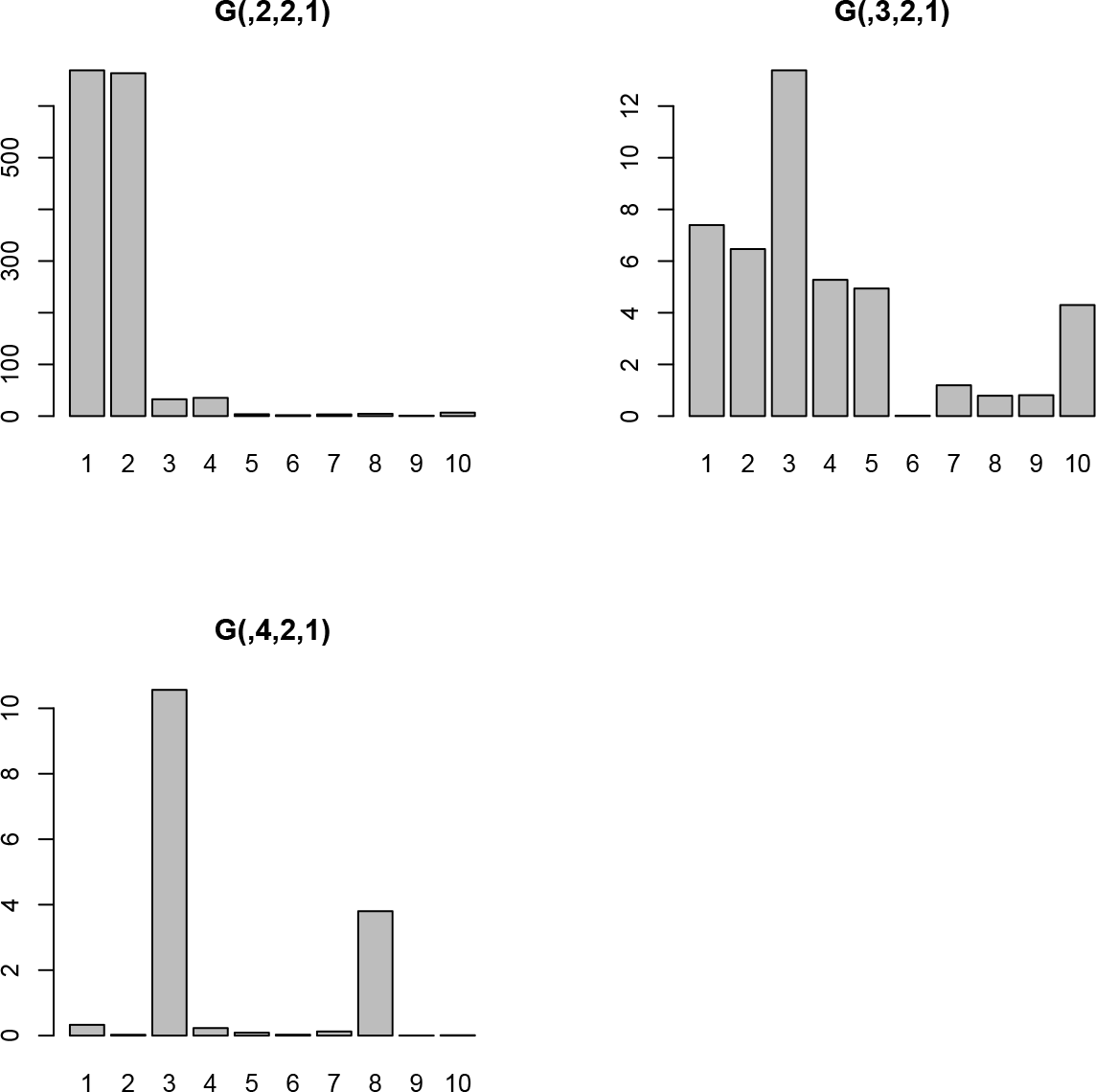
|***G*(*ℓ***_**1**_, **2, 1, *ℓ***_**4**_)|. Horizontal axes are *ℓ*_4_.

To further confirm the coincidence of analyses between with and without genome, we draw Venn diagram (Fig. 10). Considering the large numbers of contigs (*N* = 278, 167) or genes (*N* = 24, 090), this numbed of overlaps is clearly significant. Thus, we can conclude the analysis with contigs coincide with that with genome well. We also compute AUC to predict the selected 155 contigs with *P*-values attributed to genes as follows. At first, all contigs are divided to two classes, either selected 155 contigs or others. Then, *P*-values attributed to genes are re-attributed to the corresponding contigs. These *P*-values re-attributed from genes to contigs are used to discriminate contigs between selected 155 contigs and others.

**Figure 10.**
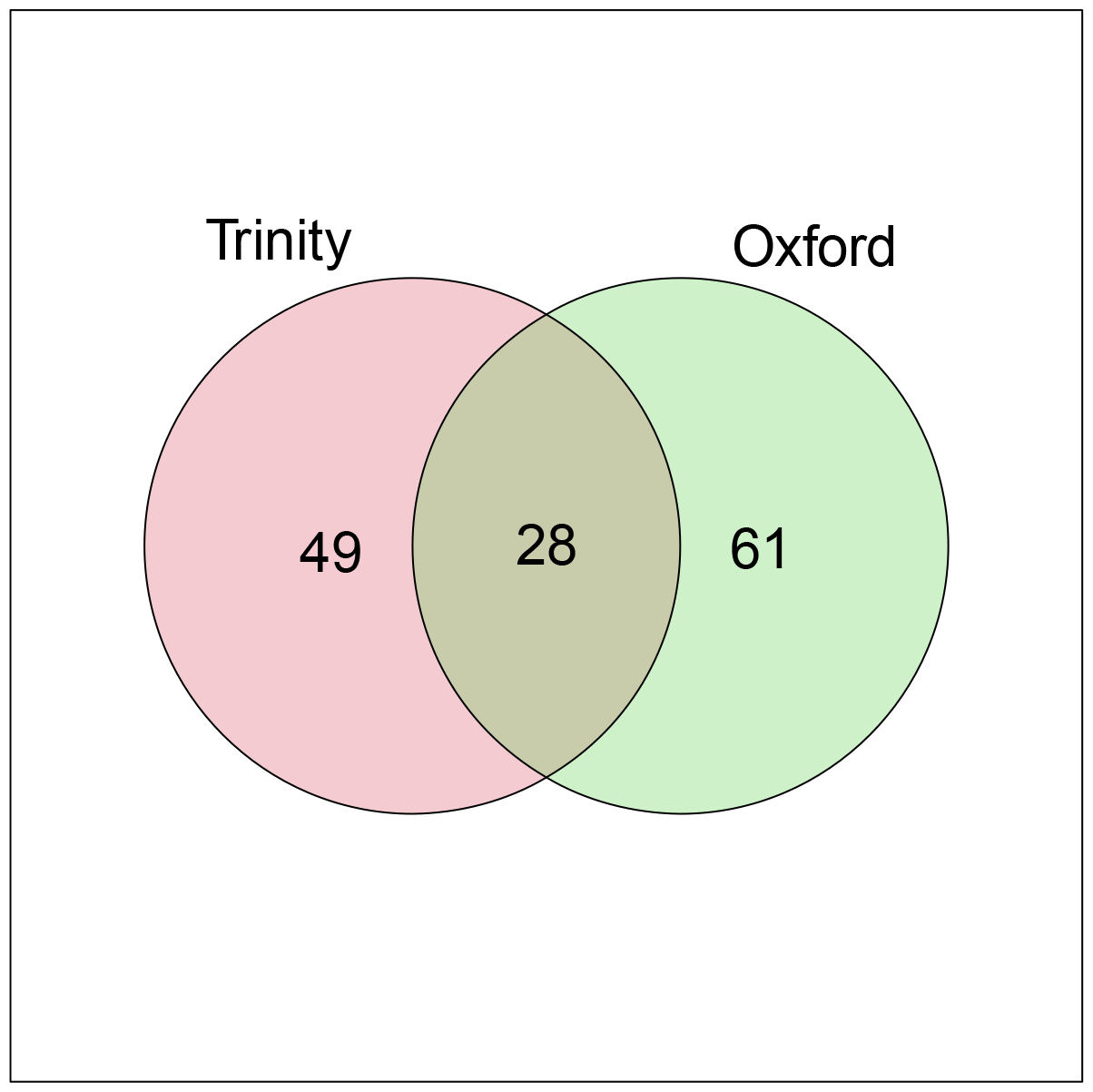
Venn diagram between the selected contigs and genes.

and found that AUC is 0.95. Thus not only selected genes/contigs, but also *P*-values themselves coincide with each other between with and without genome.

Finally, we have performed enriched analyses toward the selected 89 genes (Fig. 11). Although the number of GO terms increased from Fig. 5, there are some overlaps between Figs. 5 and 11. Thus, TD based unsupervised FE has ability to select biologically reasonable genes even without genome.

**Figure 11.**
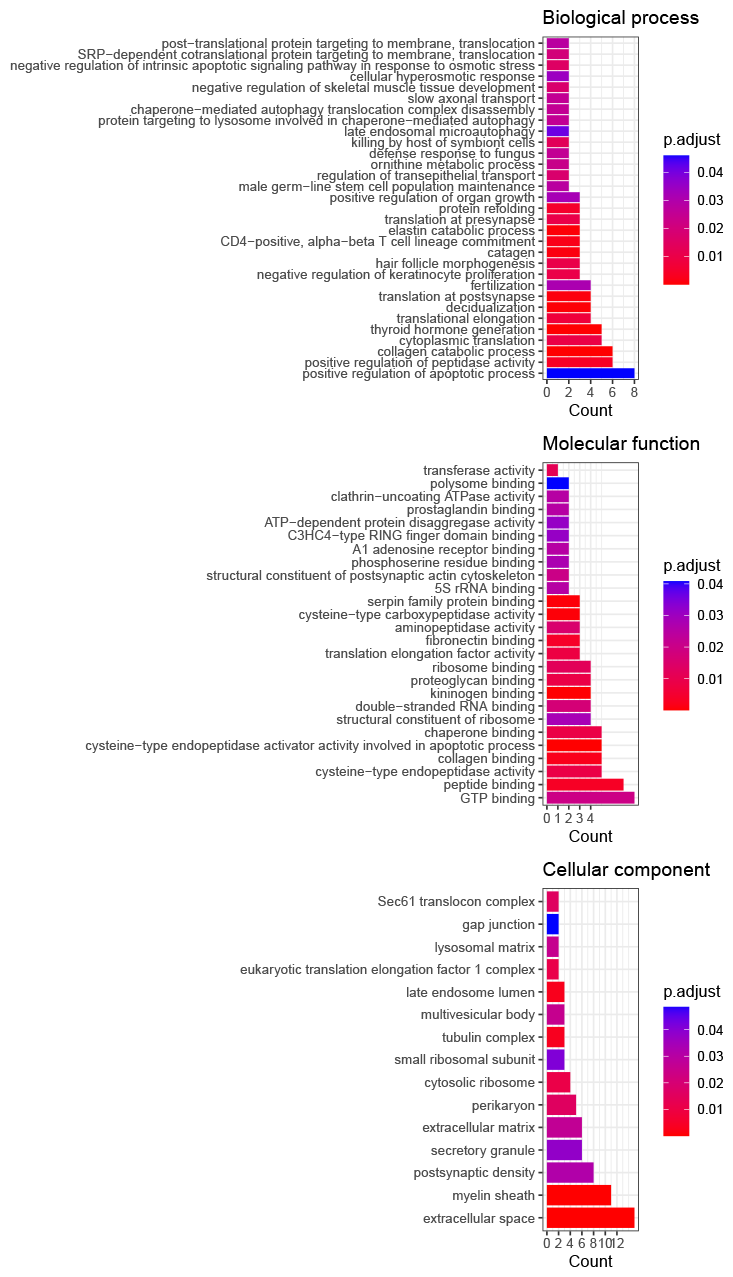
The results of GO term enrichment analysis of planarian genes selected by our method based on blast results against well annotated mouse proteins

Yet another objection might be small number of genes/contigs selected. Nevertheless, we already proposed the solution to improve the selected ones [40]. Since in the case when we need more genes/contigs selected, we can employ the strategy, the small number of genes/contigs selected is not a critical problem.

## 5. Conclusions

First, TD-based unsupervised FE could identify more contigs that are expressed distinctly between normal and defective samples, as well as being expressed in a time-dependent manner (days after RNAi) then DESeq2. GO term enrichment analysis also showed these contigs might be related to planarian regeneration.

In addition, half of the identified 155 transcripts were likely alternative spliced transcripts of a longer transcript. This supports the ability of TD-based unsupervised FE to select contigs that share similar expression patterns, since alternative spliced transcripts from longer transcripts are likely to share the same expression pattern. As a result, TD-based unsupervised FE is expected to be an effective tool to be applied to DET detection using the redundant contigs obtained by applying the *de novo* assembly of short reads from RNA-seq applied to non-model organisms that lack a well-annotated genome.

Furthermore, the results obtained with contigs coincide with those with genome and gene model; it is expected that TD based unsupervsied FE can be a promising tool to deal with contigs of organism without genome.

## Supporting information

BLAST search results toward 155 contigs (excel) and contigs to genes correspnding table (csv)

## Supplementary Materials

The following supporting information can be downloaded at: BLAST search results toward 155 contigs (excel) and contigs to genes correspnding table (csv)

## Author Contributions

M.K. and N.K. conceived and conducted the experiments, and all authors analysed the results and reviewed the manuscript.

## Data Availability Statement

All quantification results and sample information of the RNA-seq analysis were deposited as GSE174855 in the GEO repository.

## Disclaimer/Publisher’s Note

The statements, opinions and data contained in all publications are solely those of the individual author(s) and contributor(s) and not of MDPI and/or the editor(s). MDPI and/or the editor(s) disclaim responsibility for any injury to people or property resulting from any ideas, methods, instructions or products referred to in the content.

